# Synolog: A Scalable Synteny-Based Framework for Genome Architecture Characterization

**DOI:** 10.64898/2026.04.07.717040

**Authors:** Giovanni Madrigal, Julian Catchen

## Abstract

Detailing the genomic architecture across multiple organisms has been a task performed for decades. The continuing growth of genomic datasets not only serves as a resource for studying genome evolution but warrants the availability of scalable and user-friendly software for processing these datasets. Here, we present Synolog, a bioinformatic toolkit that can automatically identify orthologs for both protein-coding and non-coding genes, synteny clusters across two or more genomes, as well as retrogenes, and segmental duplications. Applying Synolog, we illustrate cases of local gene expansions in ecologically disparate turtle species, identify synteny clusters across hundreds of millions of years of metazoan evolution, and reconstruct chromosome-level assemblies in teleosts using the inferred synteny clusters; all using its integrated visual features. In parallel, we compare our orthogroup method to that of commonly used software and note the tradeoffs of making inferences solely based on sequence similarity versus a synteny-based approach.

## Introduction

The architecture of a eukaryotic genome is described by the organization and structure of a set of genomic elements – from simple sequences to information-rich, large-scale structured DNA – and the chromosomes that encapsulate these elements. This organization evolves over time yet must be stable enough to produce a functional physiology and ensure fitness (Misteli 2007; Rowley and Corces 2018). Characterizing genome architecture has proved to be useful in several biological fields. For example, by analyzing genome architecture, researchers have been able to illuminate phylogenetic relationships (Simakov et al. 2020), identify highly diverged homologs (Pegueroles et al. 2019; Yu et al. 2026), and study the evolution of gene families (Harduin-Lepers et al. 2008; Widdison and Coffey 2011; Zhang et al. 2024). Although genome architecture can be examined from different aspects, one approach for surveying the structure of two or more genomes is by identifying orthologs and examining their arrangement and structure across the focal genomes.

Orthologs are genes separated by one or more speciation events. One method for detecting orthologs is by comparing sequences (e.g., using BLAST (Altschul et al. 1990)), which allows for the identification of reciprocal best hits (RBHs) – a pair of genes that are best matched with one another based on sequence similarity. Given location information, a set of genomes can be screened for regions containing colocalized orthologs. These genomic neighborhoods that persist through evolutionary time are said to display conserved synteny. Several bioinformatic tools that employ orthologs to detect conserved synteny have been published (for a review, see Lallemand et al. 2020); however, a number of these softwares require homology relationships to be predetermined (Ludwig and Mrázek 2024; Parey et al. 2020; Zhang et al. 2025). One popular tool used for inferring orthologs is OrthoFinder2 (Emms and Kelly 2019). Under the OrthoFinder2 framework, protein-coding genes are grouped into an orthogroup based on a corrected sequence similarity score. This approach proves favorable in cases where only a single copy of a gene exists within each genome.

However, paralogs (genes arising from duplication events independent of speciation) that retain a notable amount of their sequence composition hinder ortholog inference. By considering the genetic neighborhoods a set of homologs reside in, one may illuminate the evolutionary relationship between these elements (Catchen et al. 2009). In cases where gene duplicates are found in tandem, in particular when the duplication events are recent in evolutionary time (Kono et al. 2018), discriminating between orthologs and paralogs may be challenging even under manual curation (Ye et al. 2016; Martinez Gomez et al. 2021). The ortholog conjecture suggests that in this case, the orthologs would be more similar to one another in composition and function than they would be to paralogs (Peterson et al. 2009; Studer and Robinson-Rechavi 2009; Perez et al. 2025); however, it is conceivable that paralogs may retain their ancestral function while selection is relaxed on the duplicated ortholog (Nehrt et al. 2011), or sequence homology and function may be maintained in both copies via subfunctionalization (Force et al. 1999).

In the context of conserved synteny, identifying one-to-one orthologs may not be necessary, as tandem duplicates can still be used to represent the homologous locus. For example, MCScanX (Wang et al. 2012) represents a set of tandem duplicates using the gene duplicate with the highest sequence similarity to the corresponding ortholog, while OrthoRefine (Ludwig and Mrázek 2024) permits paralogs to be merged into a synteny block if the neighboring homologs are found to be part of the same predefined hierarchical orthogroup (HOG). This flexibility allows for the construction of larger synteny blocks and permits the discovery of local gene expansions and contractions. The extent to which these benefits are realized, however, is not solely determined by the ortholog representation strategy employed.

Outside of tandemly duplicated paralogs, another type of duplication that can be inferred using sequence similarity and positional information are segmental duplications. These genomic features are duplicated, non-overlapping regions in a genome that can be thousands of base pairs in length (Bailey et al. 2001) and are considered as a major source of evolutionary novelty (Flagel and Wendel 2009; Vollger et al. 2022). For instance, genes found in segmental duplications have been reported to be involved in disease resistance (Marques-Bonet et al. 2009), cellular housekeeping (Cannon et al. 2004), and the regulation of metabolic processes (Li et al. 2024), highlighting their incorporation and maintenance over evolutionary time. Although synteny has been used to detect these genomic segments (Haas et al. 2004), more commonly used approaches involve aligning a genome or sequence to itself (Numanagić et al. 2018a; Hanke and Dagan 2025) or aligning the constituting contigs a reference genome (Dallery et al. 2017).

When gene copies occur outside of syntenic blocks, it can be difficult to differentiate paralogs from orthologs and grouping these types of genes based only on RBH information can be misleading (Watanabe et al. 2023). However, sequence similarity and gene localities can be applied to pinpoint retrogenes – duplicated genes generated through retrotransposition (Weiner et al. 1986; Brosius 1991). A notable, but not constant, feature of these types of elements is a lack of introns (Brosius 1991) and regulatory elements (Mighell et al. 2000), a poly(A) tail and adjacent repetitive sequences (Vanin 1985). These elements are often inserted in a location distant from the paralogous copy (Dai et al. 2006; Cusack and Wolfe 2007); information that has previously been implemented for retrogene identification (Carelli et al. 2016). Therefore, the spatial relationship between retrogenes and their parental copies renders conserved synteny a robust basis for retrogene identification.

As evolutionary distance (i.e., time since the last common ancestor) increases between two species, the more likely processes such as gene conversion, gene loss or gain, and small insertions or deletions can blur orthologous relationships (Dalquen et al. 2013). As such, *ad hoc* methods are routinely implemented to form orthogroups for the analysis of deep phylogenies (Zeng et al. 2017; Walden and Schranz 2023; Simakov et al. 2020; Lai et al. 2025; Alam et al. 2025). It has previously been reported that the formation of orthogroups through an RBH or similar approach could introduce phylogenetic error if the aim is to reconstruct a species tree (Koski and Golding 2001). Although some studies have used synteny to refine the predicted orthogroups, these research efforts required the use of multiple or custom softwares (Simakov et al. 2020; Walden and Schranz 2023; Alam et al. 2025), highlighting the need for a single resource for inferring orthogroups that leverages both sequence similarity and synteny.

Synteny can be used to clarify orthologous relationships amongst a set of genes between two or more distantly related taxa but can also define orthogroups for relatively fast-evolving loci between closely related taxa (Pegueroles et al. 2019). Compared to protein-coding genes, genomic elements such as non-coding RNAs (ncRNAs) have been applied to a lesser extent to identify both orthogroups and synteny blocks. This is partially due to the low sequence-conservation between homologs (Noviello et al. 2018), yet previous work has shown that ncRNA can be used to find homologs between distantly related taxa such as humans and zebrafish (*Danio rerio*) (Hezroni et al. 2015). Furthermore, not only can ncRNA represent a considerable proportion of an organism’s gene content (Wallberg et al. 2019), the plethora of biological functions these elements are predicted to exhibit (reviewed in Mattick et al. 2023) make these loci an invaluable resource for comparative genomics (Lee et al. 2019; Zhou et al. 2025).

The size of a synteny block is largely influenced by the quality of the genome assembly used (Liu et al. 2018) and the evolutionary distance between focal genomes. Intuitively, syntenic regions that are split across different genomic contigs or scaffolds are reported as separate synteny blocks. Several tools have been developed to take advantage of synteny to scaffold fragmented assemblies by using a more complete reference genome from an evolutionarily close relative to guide the scaffolding process (Aganezov et al. 2015; Aganezov and Alekseyev 2016; Bao et al. 2014; Tamazian et al. 2016; Anselmetti et al. 2018; Waterhouse et al. 2020); yet these scaffolding-centered tools depend on users applying other software to obtain syntenic information for their genomes of interest.

Long-read sequencing and chromosomal capture libraires have accelerated the number of high-quality genomes sequenced and assembled, enabling researchers to examine genome architecture across diverse and large datasets. This increase in genomic data warrants a scalable framework to process these large datasets and readily obtain results. Here, we present Synolog, an automated framework for identifying orthogroups for both protein-coding and non-coding genes, conserved synteny, segmental duplications and other genomic features. The core of Synolog is implemented in C++, with utilities provided in Python for business logic and visualization with virtually no dependencies. The system operates on a set of standard files describing the set of genomes of interest (e.g., FASTA, AGP, GTF) and provides convenient exports for further analysis (e.g., a gene counts file or orthogroup-specific FASTA files).

In this work, we demonstrate Synolog’s utility across a diverse set of taxonomic groups and illustrate its ability to generate a chromosome-level assembly using the predicted synteny clusters. We applied the Synolog pipeline cross three datasets that comprise of a wide range of evolutionary time scales, ecologies, and genome architectures. In the first case study, we examine species in the order Testudines and compare our ortholog inference algorithm to that of OrthoFinder2, while simultaneously using synteny to uncover genomic features that may be associated with ecological adaptation. In our second investigation, we again compare our ortholog inference method to that with OrthoFinder2, this time demonstrating the identification of synteny clusters in a deep phylogeny. For the last case study, we demonstrate the utility of our synteny scaffolding pipeline by generating chromosome-level sequences from contig-level genome assemblies of notothenioid fishes. These results highlight the variety of biological questions that can be addressed with Synolog while demonstrating its novelty in the field of comparative genomics.

## Results

Building upon the previous works of Catchen et al. (2009) and Small et al. (2016), we developed Synolog, a framework for synteny-based analyses to identify orthogroups, conserved synteny blocks, segmental duplications, and retrogenes. Before presenting the results of three case studies, we first describe how the software works, with details in the Methods below.

One of the barriers to genome architecture analysis is the number and type of data files required to describe it. When analyzing a small number of genomes, Synolog can be directly executed on the command line with the genome data files directly provided.

However, once an analysis reaches a certain size, Synolog provides the *species cache*, which is an organized set of data, including gene locations (GTF/GFF files), genome structure (AGP files), precomputed BLAST hits of different types, among others (Fig. 1A). This cache is managed by the `synolog_cctl.py` program and the user adds/removes/changes files for an analysis using this interface. When using the species cache, it is provided to the core Synolog program which understands the cache and where to find the appropriate data within it. Several other utilities are provided to facilitate the collection of BLAST hits for the genomes in the species cache.

**Figure 1:**
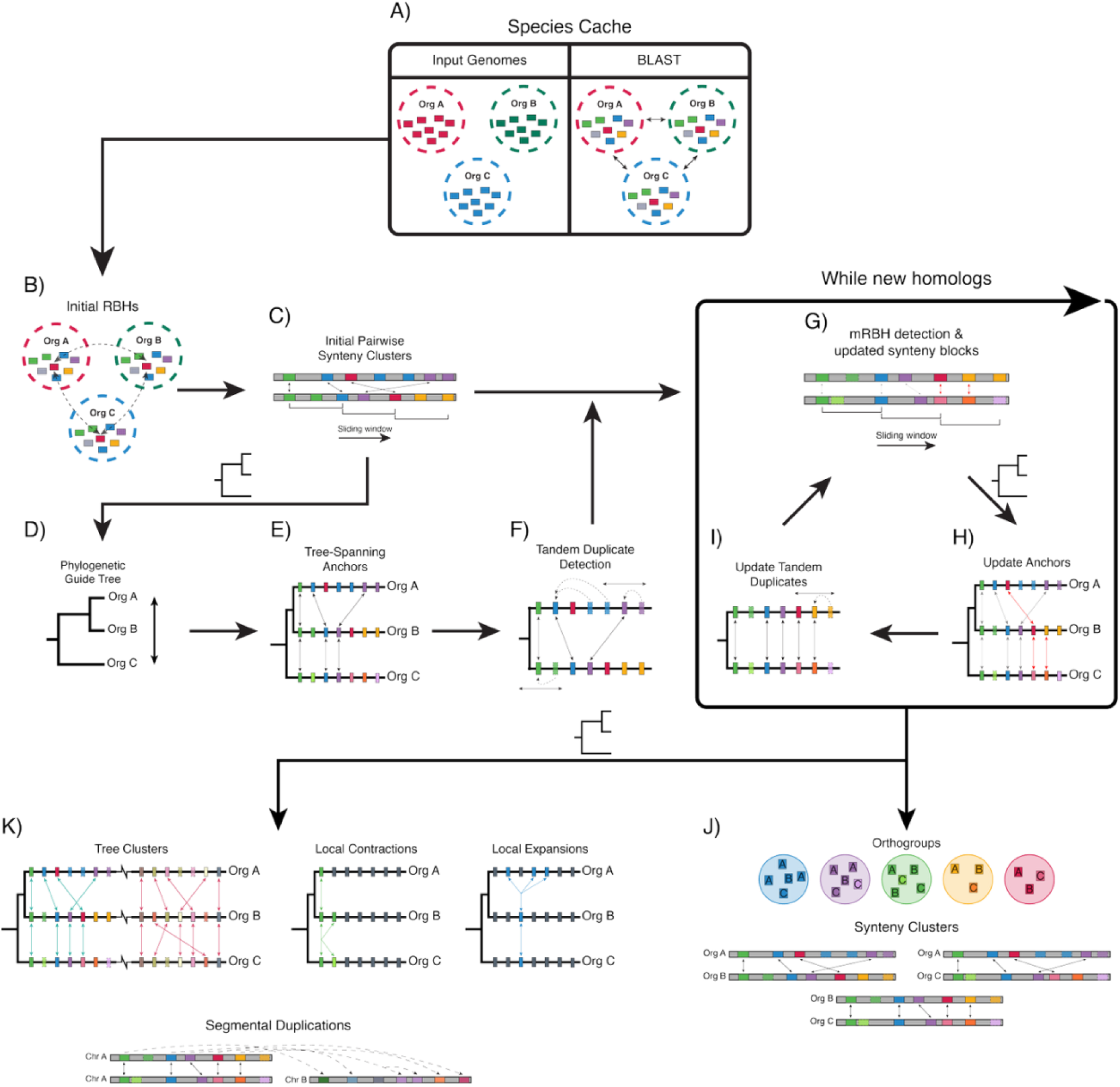
Illustration of the Synolog pipeline. (A) The structure of the genomes and the BLAST hits between genomes are stored into the species cache prior to beginning a comparative analysis. (B) The pipeline begins by inferring reciprocal best hits (RBHs) between the queried organisms prior to (C) establishing the initial synteny clusters using a sliding window technique. If a phylogenetic tree is provided, (D) the organisms are linearally organized based on the provided phylogeny, which is then used to (E) detect tree spanning anchors (syntenic orthologs across all organisms in the tree) and (F) conduct an initial tandem duplication search. (G) Genes that did not have a syntenic RBH or marked as duplicates are then subjected to continuous iterations of a modified RBH (mRBH) algorithm that further builds the inferred synteny clusters until no new homologs can be detected. If phylogenetic information is present, (H) new anchors (which may or may not extend across the tree) are determined (H) followed by (I) another tandem duplicate search. (J) Orthogroups and pairwise synteny clusters are generated once there are no updates to the orthogroups and synteny clusters. (K) Assuming that a phylogenetic tree is available for guidance, local gene contractions and expansions relative to the most basal organism are inferred, along with synteny clusters spanning three or more organisms (i.e., tree clusters) and pairwise segmental duplications.

The execution of the core Synolog program begins with the identification of all pairwise RBHs (Fig. 1B), after which the analysis employs a sliding window to construct an initial set of synteny blocks based on co-localized orthologs across these comparisons (Fig. 1C). Initially designed to work in this pairwise manner, Synolog is now capable of using phylogenetic information, when the user supplies a tree, to guide the analysis to compare the most closely related species and move towards more distantly related taxa in search of syntenic orthologs (Fig. 1D). Taking this phylogenetic tree into context, syntenic orthologs are scanned for orthogroups that concordantly span the phylogeny, creating an initial set of phylogenetic anchors (Fig. 1E). These syntenic tree-spanning phylogenetic anchors are then subjected to a tandem duplication algorithm, where a sliding window is used to identify adjacent gene duplicates for a given phylogenetic anchor using the other anchors in the current syntenic group (Fig. 1F).

Once the initial synteny clusters and orthologs are identified, Synolog applies an iterative approach in which a modified RBH (mRBH) method is used to detect additional orthologs within a shared synteny block, progressively expanding the set of synteny blocks (Fig. 1G). Following, additional phylogenetic anchors can then be constructed (Fig. 1H) and subjected to a tandem duplicate search (Fig. 1I). This process is repeated until no new information can be obtained, at which point the inferred orthogroups and synteny clusters are reported (Fig. 1J). Pairwise synteny blocks and phylogenetic information can then be leveraged together to perform a tree clustering algorithm, producing synteny clusters that span three or more species based on their shared syntenic orthologs (Fig. 1K). Additionally, each orthogroup is evaluated for local gene contractions and expansions by designating the most basal species as a reference point, enabling the identification of reduced or increased gene copy numbers across the more derived species (Fig. 1K).

Synolog also leverages its inferred pairwise synteny blocks to identify additional genomic features beyond orthologs. Segmental duplications are detected by first examining the syntenic orthologs within each synteny block to identify foreign paralogs (i.e., paralogs outside of the known synteny block). A sliding window is then employed to construct gene neighborhoods resembling the original synteny block based on these foreign paralogs. Retrogene identification is achieved through a complementary approach, whereby multi-exon orthologs are used to identify single-exon paralogs located on a distant chromosome.

To aid researchers in their analyses, we provide users with an easy-to-use Python program to immediately visualize their results (`synolog_plot.py`). We additionally developed a synteny-based scaffolding procedure that uses the inferred synteny blocks to scaffold a set of contigs into a chromosome-level assembly, enabling more robust downstream analyses (`synolog_collinearize.py`). We also provide programs to provide detailed information on the Synolog analysis for particular genes or synteny blocks (`synolog_info.py`) as well as to export sequences from the different, defined groupings described above (`synolog_fasta.py`).

### Case Study 1: Testudine Evolution

Turtles have been described as a slowly evolving lineage relative to other taxonomic groups (Shaffer et al. 2013), yet they have evolved to inhabit a diverse array of environments from the sea to the desert (Ernst et al. 1998). In a previous study, Arantes et al. (2025) revealed a high level of conserved synteny across 11 species of testudines representing five ecotypes; however, this effort was focused on sea turtles and the orthologs used were detected using a conservative RBH methodology. Here, we inferred conserved synteny across five species spanning five ecologically distinct habitats and more than 100 million years of evolution (Arantes et al. 2025). These species include the green sea turtle (*Chelonia mydas*; marine), the diamondback terrapin (*Malaclemys terrapin centrata*; estuarine-adapted), the red-eared slider (*Trachemys scripta elegans*; freshwater) the Aldabra giant tortoise (*Aldabrachelys gigantea*; terrestrial vegetative) and the Mexican gopher tortoise (*Gopherus flavomarginatus*; terrestrial desert). Although the chromosome-level genome assemblies obtained for each species were comparable in size (approximately 2.21-2.25 Gb), the haploid chromosome number varied, with three species possessing 25 chromosomes, the Aldabra giant tortoise retaining 26, and the green sea turtle at 28 (Wang et al. 2013; Jiang et al. 2025; Simison et al. 2020; Çilingir et al. 2022; http://www.vertebrategenomesproject.org/). Moreover, the total number of genes, both coding and non-coding, also differed between species, with a difference of more than 10,000 genes between the Aldabra giant tortoise and the red-eared slider (Simison et al. 2020; Çilingir et al. 2022). We processed this data through Synolog and used the phylogeny reported by Thomson et al. (2021) to guide the algorithm.

After applying Synolog, we found that the overall number of protein-coding genes grouped into an orthogroup is comparable to that reported by OrthoFinder2 (Fig. 2A). In three out of the three species, Synolog detected slightly more protein-coding orthologs than Synolog (Table S1); however, with the inclusion of non-coding genes, Synolog identifies several thousand more orthologs than OrthoFinder2 for all five species (Fig. 2A). The proportion of protein-coding genes placed into an orthogroup was over 90% for four of the five species by both methods, while Synolog assigned approximately 2,000 non-coding genes in each species (Fig. 2A). Exact gene and ortholog counts for each species can be viewed in Table S1.

**Figure 2:**
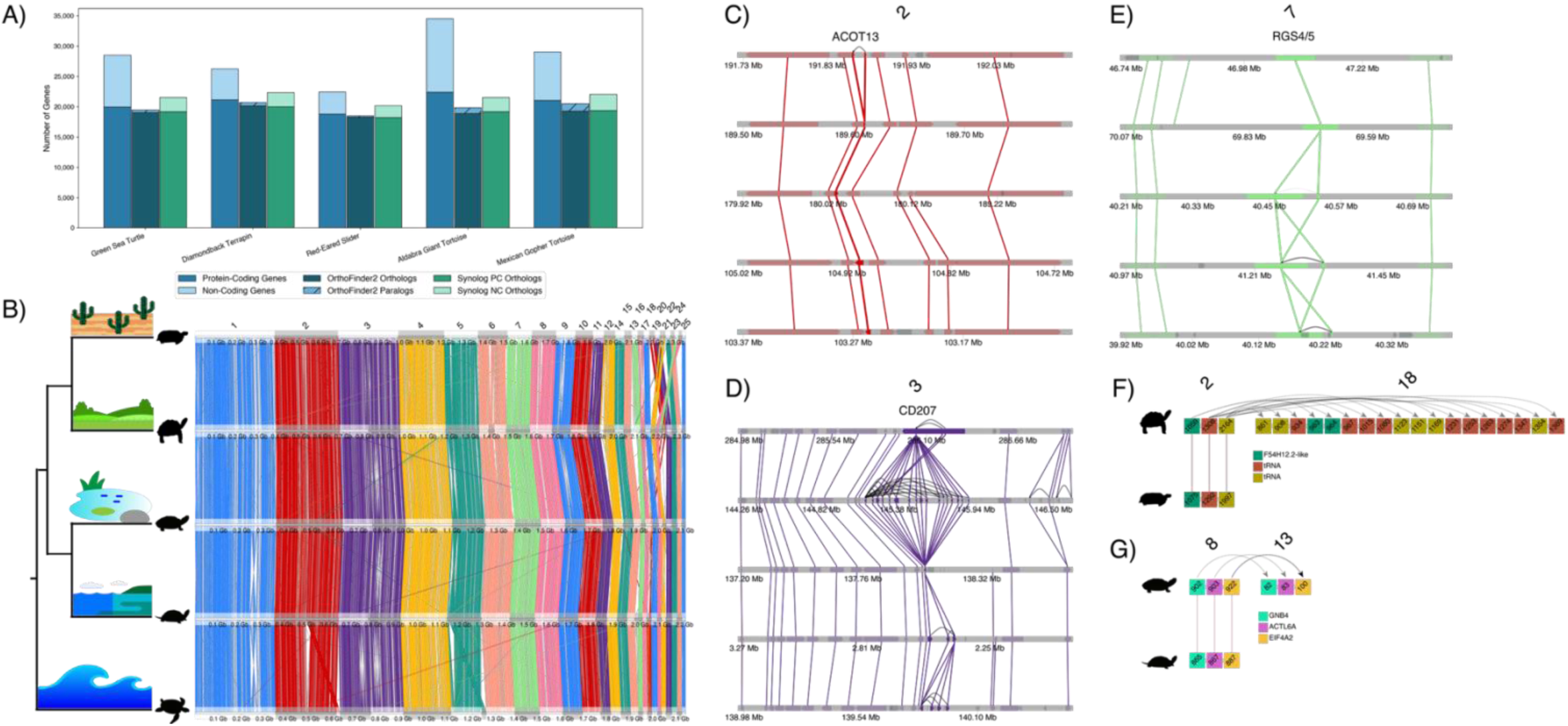
Overview of ortholog inferences and conserved synteny across testudines. (A) The number of protein-coding orthologs is comparable between OrthoFinder2 and Synolog; however, including non-coding genes yielded more orthologs in general. (B) Turtle species are arranged in the tree based on the phylogeny in Thompson et al. (2021), with the Mexican golpher tortoise (*Gopherus flavomarginatus*) at the top, followed by the Aldabra giant tortoise (*Aldabrachelys gigantea*), the red-eared slider (*Trachemys scripta elegans*), the diamondback terrapin (*Malaclemys terrapin centrata*), and the green sea turtle (*Chelonia mydas*). The phylogenetic tree is annotated with a cartoon of the type of habitat occupied by each species. Genome-wide synteny plot generated by Synolog depicts widespread conserved synteny across all species. Regional plots produced by Synolog illustrate examples of local expansions, where (C) there is a tandem duplication of ACOT13 in the Mexican gopher tortoise, (D) a large local expansion of C-type lectin genes in the Aldabra giant tortoise relative to other turtles, and (E) a difference in copy number of RGS genes where terrestrial tortoises contain a single copy while turtles that experience aquatic or marine environments have two copies. The ordering of species for the regional plots mimics the ordering in A. Outside the synteny clusters, (F) three genes, one being noted as a longevity-related gene, are segmentally duplicated and expanded across two chromosomes in the Aldabra giant tortoise relative to the Mexican tortoise. In another case, (G) three segmentally duplicated genes, two of which have been documented to be involved in environmental adaptation, are detected in the diamondback terrapin when compared to the red-eared slider.

Using the `synolog_plot.py` subprogram, it is apparent that conserved synteny is largely kept intact across the genomes despite the karyotype and ecological differences between the species (Fig. 2B). Synolog’s automatic tree cluster inference algorithm identified 916 tree clusters, of which 208 clusters contained genes from all five species and with 27 consisting of at least 1,000 orthologs across the five species. Although these results align with those reported by Arantes et al. (2025), Synolog generated them without intervention and produced visualizations without relying on third-party software.

Synolog reported a total of 19,862 orthogroups containing 107,667 orthologs across the five turtle genomes. Of these 19,862 orthogroups, 12,201 (61.42%) were identified as one-to-one orthogroups, i.e., one ortholog from each of the five species. OrthoFinder2 reported 99,219 protein-coding genes encompassed in 17,990 orthogroups. Of these 17,990 orthogroups, 361 orthogroups containing 3,472 genes were flagged as species-specific – emphasizing that OrthoFinder2 will report groups of orthologs that consist of genes entirely from one species (Fig. 2A). Removing these paralogous gene groups reduced the orthogroups to 17,629 and orthologs to 95,747. Interestingly, 2,150 of these reported paralogs were placed in an orthogroup by Synolog. To illustrate an example, OrthoFinder2 placed two *LTA4H*-like genes in the red-eared slider into a single paralogous group; however, Synolog was able to classify these two genes as tandem duplicated genes to the adjacent *LTA4H* gene and merge them into the respective orthogroup (Fig. S1). Similar circumstances likely resulted in OrthoFinder2 reporting 261 more single-copy orthogroups than Synolog.

Synolog is not limited to protein-coding genes (in contrast to OrthoFinder2) and can also examine non-coding genes. In total, Synolog reported 11,725 non-coding genes compiled into 2,905 orthogroups. Many of these genes are syntenic and found as tandem duplicates. As an example, we observed a case on chromosome 2 where the *CCM2* and *NACAD*-like gene are flanked by syntenic non-coding genes (Fig. S2). Along this chromosome, we also identified a locus containing variable numbers of non-coding homologs between all five species, including a lineage-specific loss of non-coding elements in the red-eared slider (Fig. S3), Lastly, Synolog’s tree clustering algorithm identified 2,881 tree-spanning non-coding orthologs embedded alongside 60 tree-spanning orthogroups Amongst the orthologs reported, Synolog labeled 493 genes as candidate retrogenes. The number of predicted retrogenes varied across the turtle species; 184 retrogenes for the Aldabra giant tortoise, 113 for the diamondback terrapin, 82 for the Mexican gopher tortoise, 78 for the green sea turtle, and 36 for the red-eared slider. To note an example, Synolog was able to identify three interferon retrogenes in the Mexican gopher tortoise, a gene family that has been reported to evolve via retrotransposition in primates (Schelle et al. 2023). OrthoFinder2 grouped these three genes within the same paralogous orthogroup but was unable to identify their retrogene nature.

We further sought to associate genomic features with environmental adaptation by identifying increased gene copy numbers for potentially ecologically relevant genes. In our analyses, we found 20 orthogroups where strictly terrestrial turtles had a higher copy number than their counterparts and 41 orthogroups for which higher copy number was observed in strictly aquatic species. Additionally, a total of 40 orthogroups had higher copy numbers for species adapted to saline waters. Considering turtle species individually, we found 77, 95, and 796 orthogroups in which the Mexican gopher tortoise, the green sea turtle, and Aldabra giant tortoise had the highest gene counts, respectively, representing species-specific local gene expansions.

To further ground Synolog’s results with turtle biology, we performed a gene ontology (GO) annotation analysis on the orthogroups identified and found a number of biological processes that are consistent with environmental adaptation. For example, the GO terms water transport (GO:0006833), urea transmembrane transport (GO:0071918), regulation of heart rate by cardiac conduction (GO:0086091), and embryonic skeletal system development (GO:0048706) were found in orthogroups that contain more gene copies in species inhabiting aquatic habitats compared to the strictly terrestrial species. For species adapted to terrestrial environments, we found fewer enriched GO terms, with examples such as response to regulation of cytokine production (GO:0001817) and positive regulation of smooth muscle cell proliferation (GO:0048661). For the green sea turtle and the diamondback terrapin, two species that experience saline conditions, we found orthogroups linked to biological processes such as regulation of protein folding (GO:0006457) and the G protein-coupled receptor signaling pathway (GO:0007186), which may play a role in maintaining protein stability and increased tolerance under salinity stress (Misra et al. 2007; Leprêtre et al. 2025).

To note specific genes that may be involved in ecological adaptation, we present a case with *ACOT13*, which was found to be tandemly duplicated exclusively in the Mexican gopher tortoise (Fig. 2C). Copy number variation of this gene has previously been associated with desert adaptation (Chebii et al. 2021), which may aid in fat storage, an advantageous trait given the sparse nutritional resources found in desert environments (Olsen et al. 2021). To illustrate a second case, we found 16 tandem duplicates of C-type lectin domain family genes in chromosome 5 of the Aldabra giant tortoise (Fig. 2D). Although it is unclear whether these genes may have played a role in environmental adaptation, C-type lectin genes are recognized as a marker for Kupffer cells in mouse models (Yang et al. 2013), which are cells involved in the liver’s immune response.

Experimental validation conducted by Jiang et al. (2021) showed that *CLEC4F* is essential for Kupffer cells to clear desialyated platelets during a bacterial infection. In another example, *RGS5* is found as a single copy in two tortoise species but tandemly duplicated in the other turtles and annotated as *RGS4* (Fig. 2E). Interestingly, previous work has reported changes in gene expression of *RGS4* in response to different salinity treatments in a semi-anadromous fish (Komoroske et al. 2016), while *RGS5* has been noted to increase in expression in hypoxic conditions (Jin et al. 2009).

Lastly, we inspected the reported pairwise segmental duplications inferred by Synolog for possible ecologically relevant genes. In one example, when comparing the Aldabra giant tortoise to the Mexican gopher tortoise, we found a segmental duplication in the Aldabra giant tortoise containing two copies of an uncharacterized protein resembling *F54H12*, a gene described as a longevity gene in *Caenorhabditis elegans* (Hamilton et al. 2005) (Fig. 2F). The remaining genes contained in this segmental duplication were non-coding genes. In another instance, we found a segmental duplication in the diamondback terrapin harboring *EIF4A1*, a gene that has been reported to influence salt tolerance in rice (Chen et al. 2025), and *GNB2*, a gene highlighted by Marques et al. (2021) when investigating the genomic differences between marine and freshwater threespine stickleback (*Gasterosteus aculeatus*) (Fig. 2G).

### Case Study 2: Metazoan Evolution

Motivated by the application of synteny to analyze genome architecture across very distantly related taxa, we sought to replicate the work of Simakov et al. (2022), who used synteny to study genome evolution spanning more than 600 million years of metazoan history (Yin et al. 2015). However, instead of recapitulating their bespoke analysis, we wanted to see how our automated approach would compare. We applied Synolog to infer orthologs and conserved synteny on the same distantly related species used by Simakov et al. (2022): the Florida lancelet (*Branchiostoma floridae*), the flame jellyfish (*Rhopilema esculentum*), the Yesso scallop (*Patinopecten yessoensis*), the Mueller’s freshwater sponge (*Ephydatia muelleri*), and the freshwater polyp (*Hydra vulgaris*). Expectedly, there is a considerable amount of variation in genome architecture between these organisms. For example, the flame jellyfish possess a genome size estimated at 275 Mb and haploid chromosome number of 21 (Li et al. 2020); whereas the Yesso scallop has a genome size at roughly 1.43 Gb and a haploid chromosome number of n = 19 (Wang et al. 2017). Similarly, the number of protein-coding genes also considerably differs between these five species. The ∼326 Mb genome for Mueller’s freshwater sponge is predicted to have over 39,000 protein-coding genes while the smaller genome of the flame jellyfish is reported to harbor less than 15,000 protein-coding genes (Kenny et al. 2020; Li et al. 2020).

Considering that these species have not shared a common ancestor for over 600 million years (Yin et al. 2015), we ran Synolog with a relaxed sequence similarity threshold (defined in the Methods) during tandem duplication assignments. Synolog identified 53,035 orthologs organized into 11,172 orthogroups. For each species, OrthoFinder2 reported more orthologs than Synolog; however, we found many of the reported genes to be encompassed in species-specific groups (Fig. 3A). Excluding the single-species orthogroups, OrthoFinder2 reported 63,265 orthologs within 10,424 orthogroups, 2,197 of which were single-copy orthogroups. In the work of Simakov et al. (2022), a total of 2,361 mutual best hits (MBHs) were used to reconstruct conserved, ancient linkage groups (ALGs). In our analysis, Synolog identified 2,325 single-copy orthogroups. The number of predicted orthologs from both methods can be seen in Table S2.

**Figure 3:**
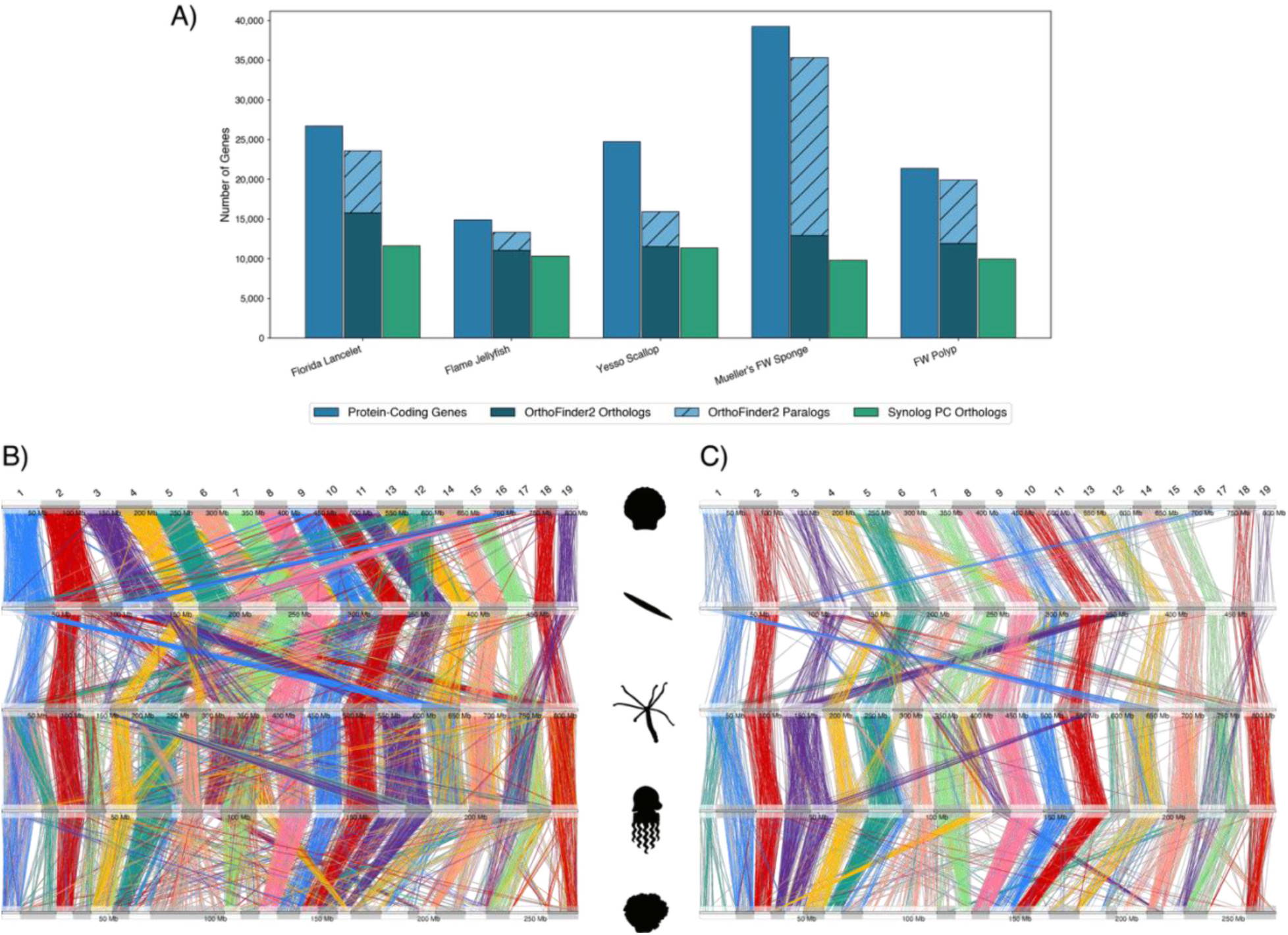
Ortholog inference and automatic detection of conserved synteny across metazoans. (A) Ortholog inference for protein-coding genes by Synolog and OrthoFinder2. (B) Synolog is able to identify conserved synteny between distantly related metazoans. In B, the Yasso scallop (*Patinopecten yessoensis*) is shown at the top of the synteny plot, followed by the Florida Lancelet (*Branchiostoma floridae*), the freshwater polyp (*Hydra vulgaris*), the flame jellyfish (*Rhopilema esculentum*), and the freshwater sponge (*Ephydatia muelleri*) at the bottom. (C) Synolog illustration that depicts the subset of orthologs that syntenically agree across all species.

When visualizing all the syntenic relationships inferred, the presence of ALGs is masked by the presence of non-collinear orthologs and lineage-specific losses (Fig. 3B); however, these genomic elements become more apparent when restricting the view to syntenic genes spanning the phylogeny (Fig. 3C), which are natively produced by Synolog. Our tree cluster algorithm identified 385 tree clusters containing syntenic genes from all five species. The reported tree clusters were then further processed to identify ALGs by selecting only those containing genes across all five species and merging them based on the encompassing chromosomes. This resulted in 35 putative ALGs that contained 3,525 genes. Upon removing ALG clusters that contained less than two genes per species (i.e., clusters not containing syntenic co-localized elements), the number of candidate ALGs reduces to 26 clusters harboring 3,480 genes. Simakov et al. (2022) reported 29 ALGs, which were composed of 2,361 MBH clusters of orthologs. In agreement with the original analysis, our analysis found that some ALGs are isolated to single chromosomes while other ALGs share chromosomal assignments within a single species (Simakov et al. 2022). While minor differences in the ALGs are observed relative to Simakov et al. (2022), these likely reflect the distinction between their manual curation approach and the automated pipeline employed by Synolog.

### Case Study 3: Synteny-Based Scaffolding

To further leverage the syntenic information discovered by Synolog, we developed a genome scaffolding approach that uses the inferred synteny clusters to generate chromosome-level genome assemblies. With the increasing number of publicly available chromosome-level reference genomes, researchers can use these resources to guide the scaffolding processing of a more fragmented assembly. Consequently, by feeding Synolog the synteny clusters between two species, with one to serve as the guide or template assembly and the other as the guided genome assembly, Synolog can identify the orientation and placement of sequences within the guided assembly to maximize the collinearity between the two genomes. This arrangement can be determined by employing the `synolog_collinearize.py` while the contigs and/or scaffolds can be physically fused using the `alter_genome_structure.py` utility.

For a proof of concept, we sought to reconstruct chromosome-level assemblies for the mackerel icefish (*Champsocephalus gunnari*) and the Antarctic bald notothen (*Trematomus borchgrevinki*) using only contig-level assemblies and synteny information. As members of the Antarctic notothenioid radiation, these two species have shown high levels of conserved synteny with the Patagonian blennie (*Eleginops maclovinus*) (Rivera-Colón et al. 2023; Cheng et al. 2023; Rayamajhi et al. 2025), a basal non-Antarctic notothenioid, despite not sharing a common ancestor for tens of millions of years (Bista et al. 2023). Consequently, we chose to use *E. maclovinus* as a template for our scaffolding approach.

Using the chromosome-level assembly of *E. maclovinus* (Cheng et al. 2023) and the phylogeny predicted by Near et al. (2018), we performed an initial Synolog analysis with all three species to generate pairwise synteny clusters. The `synolog_collinearize.py` script was used to automatically calculate collinearity between the *E. maclovinus* reference genome and either the *C. gunnari* or *T. borchgrevinki* contig-level assemblies, respectively. The script uses a linear regression algorithm to order contigs into scaffolds by permuting contig order and continually recalculating an R^2^ value describing how tightly orthologous genes are ordered between the two genomes. The highest R^2^ values reported were 0.964 and 0.958 for *C. gunnari* and *T. borchgrevinki* respectively (Fig. 4A and 4B). The contig-level assemblies were then scaffolded using `alter_genome_structure.py` (Fig. 4C). The new assemblies showed one-to-one homology with the chromosomal sequences in *E. maclovinus* (Fig. 4D and 4E) after rerunning the Synolog pipeline with the newly fused set of scaffolds.

**Figure 4:**
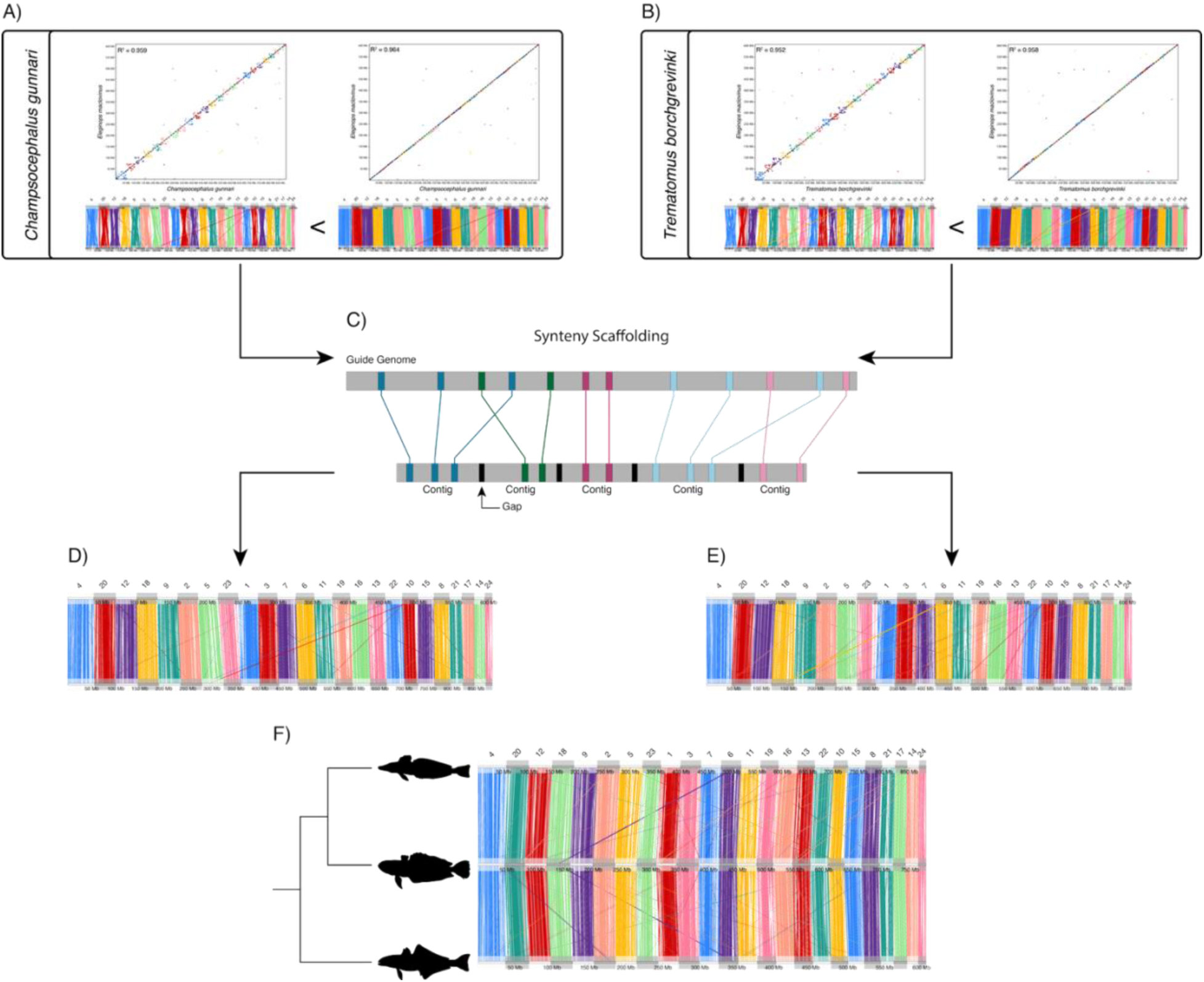
An overview of the synteny scaffolding process in notothenioids. The contig-level assemblies for the mackerel icefish (*Champsocephalus gunnari*) and the Antarctic bald notothen (*Trematomous borchgrevinki*) were processed through the synteny scaffolding procedure. Synolog examines different orderings of contigs by estimating collinearity through a linear regression using the synteny blocks. (A) In one arrangement between the Patagonia blennie (*Eleginops maclovinus*) and the mackerel icefish, Synolog preferred the arrangement with an R^2^ of 0.964 over an arrangement with an R^2^ of 0.959. (B) Similarly, Synolog selected an ordering with an R^2^ of 0.958 over an arrangement with an R^2^ of 0.952 when comparing the Patagonia blennie with the Antactic bald notothen. (C) The selected arrangements were both used to scaffold the contigs using the syntenic information. (D, E) Both the new assemblies show one-to-one homology with Patagonia blennie (*Eleginops maclovinus*) chromosome-level assembly. (F) A three-way synteny plot shows the mackerel icefish is the topmost genome, followed by the Antarctic bald notothen, and then the Patagonia blennie as the outgroup.

These results presented similar findings to the genome-wide synteny results from Rivera-Colón et al. (2023) and Rayamajhi et al. (2025), respectively (Fig. 4F).

The L50 and N50 metrics of the synteny-scaffolded genomes were measured at 12 and 39.01 Megabases (Mb) for *C. gunnari* and 13 and 33.77 Mb for *T. borchgrevinki*, reflecting strong concordance with the original Hi-C-based assemblies (Table S3). For both synteny-based assemblies, 24 sequences were designated as chromosomes based on the homologous chromosomes found in *E. maclovinus* (Fig. 4D and 4E; but see Discussion). The percentage of base pairs incorporated into these chromosomal sequences is limited by the presence of orthologous genes and differed between the two species, with the chromosome-level sequences containing 90% of the reference genome for *C. gunnari* and ∼83% for *T. borchgrevinki*.

## Discussion

Ortholog inference and synteny detection are central components in the field of evolutionary genomics. In this work, we presented Synolog and illustrated its versatility in several cases involving taxonomical groups of varying evolutionary distances. The incorporation of phylogenetic information to guide a synteny-based homology search to construct orthogroups alleviates the need to compare a genome to itself in order to identify gene duplicates. Furthermore, by including non-coding RNAs and certain types of paralogs (e.g., retrogenes), Synolog is capable of identifying conserved synteny using overlooked genomic features and provide a higher resolution of chromosomal homology across distantly related species.

Large, highly conserved syntenic regions amongst turtles has been previously identified by Arantes et al. (2025); however, by limiting their application to single-copy orthogroups, many interesting genomic features were left underexplored in their findings. By setting the most basal organism on a phylogenetic tree as a reference point, we can detect and label changes in gene copy number. Using ecologically disparate species, Synolog identified several candidate genes for ecological adaptation as either tandem duplicates or segmental duplications. Although OrthoFinder2 was able to find more protein-coding orthologs when considering all turtle species (Table S1), these genes are likely duplicated paralogs that reside outside the syntenic region containing its progenitor. Compared to OrthoFinder2, Synolog’s ortholog inference strategy is more conservative given its prioritization on synteny. As such, Synolog’s ortholog inference method is tailored towards identifying homologous regions and highlighting changes in genome architecture instead of clustering related homologs. Furthermore, syntenic non-coding orthologs were automatically detected in our analysis, further capturing the full landscape of conserved genomic features. These results illustrate that syntenic non-coding genes can be readily detectable across large evolutionary timescales despite their rapid molecular evolution, highlighting their underutilized potential in comparative genomic analyses.

While a basal organism is not equivalent to an ancestral species, we note our local expansions and contractions as relative to the most basal organism in the focal orthogroup; an approach that has previously been shown to uncover evolutionary patterns (Kim et al. 2020; Hu et al. 2022; Kim et al. 2023; Melchionna et al. 2024). Additionally, other bioinformatics tools have employed a phylogenetic tree to serve as a guide tree (Kloss-Brandstätter et al. 2011; Roosaare et al. 2017; Balaban et al. 2019), corroborating the implementation of a guide tree in our framework.

The results of Simakov et al. (2022) yielded enlightening insights into the evolution of metazoans. In this report, we were able to reproduce similar findings through an automated framework. Synolog recognized more single-copy orthogroups than OrthoFinder2 between these distantly related metazoans despite failing to group more orthologs overall; likely due to our practice of avoiding the incorporation of non-syntenic paralogs into an orthogroup apart from candidate retrogenes. The ALGs inferred from the resulting tree clusters were nearly identical in number to that found by Simakov et al. (2022), with Synolog identifying three fewer ALGs. Though further analysis is required to elucidate the evolutionary histories of these ALGs, researchers interested in studying these genomic features can use Synolog to generate results as the basis for their investigations.

In our third case study, we took advantage of syntenic information to scaffold a set of contigs into chromosome-level sequences, a procedure that also has been implemented elsewhere (Waterhouse et al. 2020). The estimated collinearity between the two notothenioid genome assemblies with *E. maclovinus* was high (> 0.95 R^2^) with both newly scaffolded assemblies containing most of the sequenced base pairs in chromosomes, indicating an algorithmic success. Though several contigs were unable to be incorporated into chromosomes due to the lack of annotated genes, this approach is a fast and cost-effective methodology to generate a chromosome-scale assembly in the absence of additional sequence data for the focal sample. However, this approach is not without limitations, as synteny-guided scaffolding is sensitive to the evolutionary distances between guide and focal genomes. It is thus important to select a guide genome that shares a similar genome architecture as the organism in question. For example, the initial assembly of *T. borchgrevinki* was shown to have a chromosomal fusion relative to *C. gunnari* and *E. maclovinus* (Rayamajhi et al. 2025), which resulted in 23 chromosomal sequences as opposed to 24. Consequently, this assembly remains incomplete, as the fused chromosome remains split in our approach. In cases like these and in absence of additional sequence data, bioinformatic approaches that leverage the initial raw reads to identify genome misassemblies exist (Madrigal et al. 2025), providing a mechanism towards a more complete reference genome.

The application of Synolog is especially useful when generating new genome assemblies. It allows one to use conserved synteny to compare the accuracy of a new, draft assembly to earlier drafts, or to other assemblies, with the capability to manually make changes to those scaffold orders. The species cache and other tools provided with Synolog make this type of application relatively easy and was applied in several notothenioid genomes (Rivera-Colón et al. 2023; Rayamajhi et al. 2025).

Altogether, we present Synolog and illustrate its ability to infer orthologs at both the protein and nucleotide level, construct synteny blocks both pairwise and across the three or more organisms as guided with a phylogenetic tree, and leverage syntenic information to identify segmental duplications, retrogenes, and scaffold a queried genome. With the growing number of genome assemblies becoming readily available, user-friendly tools for large-scale comparative genomic studies opens the door for researchers to conduct such studies and identify novel genomic insights without prior knowledge of the underlying architecture. At the time of writing, Synolog is designed to work on genes (coding and non-coding); however, this framework sets the basis for additional genome markers (e.g., transposable elements) to be used in future iterations. As such, Synolog will prove useful to address questions regarding genome architecture and provide researchers a user-friendly approach for detailing their findings.

The core of Synolog is implemented and parallelized in C++ with three sets of utility programs provided in Python. The first aids the user in organizing genome data, including genome annotations (GTF/GFF), genome structure (AGP), and gene models (FASTA) in a common location and executing programs such as BLAST. The second set of programs exports data in useful formats and aids in the scaffolding process. The final set of utilities provides a robust set of visualization tools which can aid the user in understanding the produced data as well as creating publication-level images. Synolog is provided as open-source software can be downloaded from http://catchenlab.life.illinois.edu/synolog/.

## Methods

To facilitate automation, Synolog is composed of three main components: the creation of a species cache, an orthogroups and conserved synteny analysis, and a post-inference toolkit. To illustrate Synolog’s versatility, we apply Synolog on three different case studies while simultaneously comparing the resulting orthogroups to that of the commonly used software OrthoFinder2 for the first two investigations.

### Creating The Species Cache

The species cache serves to house the genomic data required to determine orthogroups and synteny blocks. This cache is simply an ordered set of directories on disk containing all the genome-affiliated files needed by Synolog for each stage of its analysis. For example, this includes GTF or GFF3 annotation files, FASTA gene model files, and sets of BLAST hits, among others. It is initialized and interacted with through Synolog’s utility `synolog_cctl.py` and once populated can be used for multiple Synolog analyses containing different species subsets. The locations of genes along with their sequences are required entries in the cache for any given organism. Given that protein-coding genes can be represented in amino-acids while ncRNA sequences are managed at the nucleotide level, these sequences are kept separate in the cache and only one dataset is required to include an organism in an analysis. In cases where ncRNAs are annotated but not contained in its own FASTA file, users can execute the `synolog_ncgenes.py` script to extract these elements out of the reference genome so that they can be added to the cache.

Considering that a species can be represented with differing genome assemblies and gene annotations, Synolog labels each organism by a provided annotation and integration ID. Thus, multiple versions of a genome assembly/annotation set can be incorporated into a synteny analysis. To streamline the sequence comparisons amongst the integrated dataset, the `synolog_blastctl.py` script is used to launch and store a BLAST analysis for each pairwise combination for the queried organisms or species.

Synolog separates genes based on their non-coding vs protein-coding status in the BLAST stage. Non-coding RNA has been noted to evolve at a higher rate than protein-coding genes at the sequence level (reviewed in Bussotti et al. 2013). Previous efforts tailored towards non-coding RNA homology detection have employed BLAST with a shortened word_size parameter to accommodate this biology (Tang et al. 2017; Kern et al. 2018; Lemos et al. 2020; Fu et al. 2025). Accordingly, Synolog executes a BLAST search using an adjustable, shortened word_size for non-coding RNA.

### Orthogroup And Conserved Synteny Analysis

The primary analysis of Synolog is divided into two pipelines, both executed by the parallelized, C++-based, core executable `synolog`. The first pipeline identifies orthogroups and conserved synteny blocks in a pairwise manner across all the queried organisms. The second pipeline extends this approach by leveraging phylogenetic information to detect other genomic features such as tandem duplicates, retrogenes, and segmental duplications. We first explain the algorithmic approach performed in both pipelines to infer orthogroups and conserved synteny blocks before noting the discrepancies in the pipelines. A summary of the Synolog pipeline is shown in Figure 1.

### I. BLAST Processing

For the initial phase, Synolog loads the genome structure prior to processing the BLAST hits from the species cache. A percent identity of 30% or more between protein sequences has been noted to hold merit when it comes to classifying homologs (Rost 1999). Knowingly, Synolog will only retain BLAST hits with a percent identity of 30% or higher; however, in cases where no BLAST hits for a given gene meet this criterion, all BLAST hits for that gene are used. We then follow a similar approach as Catchen et al. (2009) by reducing the BLAST hits between any two queries to the singular best hit based on e-value or bitscore (user defined) and then ranking these hits. If the ranking scheme follows the e-value scheme, BLAST hits with a lower e-value are ranked higher than those with greater values. The ranking approach for bitscore runs counter to that of the e-value approach, with BLAST hits with lower bitscores being lowly ranked. Here, we note that unlike other methods, Synolog includes all gene transcripts in its analysis, and does not reduce BLAST hits for multi-transcripts genes to a set of BLAST hits from a singular representative transcript (usually the longest). This approach diverges from previous methods that choose to either use a singular transcript based on its length (Emms and Kelly 2015) or the number of BLAST hits to the other queried organisms (Altenhoff et al. 2021).

### II. Conserved Synteny Block Construction

After processing the BLAST hits, gene pairs between two organisms exhibiting a RBH relationship are labeled as orthologs. The sliding window approach described in Catchen et al. (2009) is then used to establish the initial synteny blocks. Briefly, a sliding window of size *N* genes (default 100, but user modifiable) is used to find regions between the two focal genomes where co-localized orthologs no greater than *N* genes apart can be grouped into a synteny block. This window is extended *N* genes for every additional incorporation and is implemented in the forward and reverse direction to account for orthologs that are in reverse orientations in one of the two organisms due to chromosomal rearrangements. Synteny blocks are then merged if the distance between the two blocks does not exceed *N* genes.

Once syntenic information has been obtained, the analysis shifts to detect orthologs that do not exhibit a RBH relationship. For each unmatched gene in the query organism, a modified RBH (mRBH) approach is performed where BLAST hits are processed from the highest ranked to the lowest ranked for genes in the subject organism that lack an ortholog to the query organism. Both unmatched genes are evaluated to determine if they occupy a shared syntenic region between the query and subject organisms. If the two genes are shown to inhabit the same synteny block, they are marked as orthologs. After applying this mRBH approach, the synteny block algorithm is repeated to produce larger (i.e., more gene rich) synteny blocks. With an updated set of synteny blocks, the search for orthologs using the mRBH methodology and reconstruction of synteny blocks is continuously repeated until no orthologous pairings can be found to expand the synteny blocks. The pairwise synteny blocks and ortholog inferences are then reported to the user. After all the pairwise comparisons have been concluded, orthologs across all the queried organisms are merged to form orthogroups.

### Pipeline 2: Applying A Guide Tree

To enact the additional functionalities in Synolog (e.g., tandem duplication detection), a phylogenetic tree containing the species of interest is required (in Newick format) to guide the directionality of comparisons. The organisms are ordered linearly into a guide tree based on their position on the phylogenetic tree, where the outgroup is placed at the bottom of guide tree and the topmost position is assigned to the organism occupying the most nested clade. When two or more organisms are found within this clade, the assignment is dictated by the order in the provided phylogenetic tree. Once a tree is supplied, different subsets of species can be analyzed and Synolog will prune the tree for the execution accordingly. In the beginning, this pipeline identifies orthologs exhibiting a RBH relationship and an initial set of synteny blocks identically to pipeline 1.

### III. Metric Score Correction

Unlike the independent pairwise comparisons conducted in pipeline 1, the initial orthologs are evaluated for two properties: a set of orthologs (i.e., an orthogroup) that are able to unambiguously span the phylogenetic tree while remaining syntenic with one another. These syntenic tree-spanning orthogroups are then used to compute a new scoring metric following the same logic as Emms and Kelly (2015) to correct for biased bitscores due to sequence length. For each pairwise comparison, we apply the length normalization procedure in Emms and Kelly (2015) to calculate a linear regression using only the tree-spanning orthologs. The slope and intercept are then used to correct every BLAST hit in the analysis and each gene then has its BLAST hits re-ranked based on this corrected metric. A standard deviation for each pairwise comparison is then determined using the corrected estimates. For ncRNAs, we apply the same formula but replace protein length with exonic length and keep the linear regression and standard deviation calculations separate from that of protein-coding genes.

### IV. Ortholog Refinement and Phylogenetic Anchor Construction

Given that the initial set of orthologs (based on RBHs alone) was determined without synteny or a corrected metric, paired orthologs that lack synteny are recorded and undone for re-assignment. Given that syntenic homologs are preferred over non-syntenic paralogs, we store these non-syntenic orthologs as backup orthologs to one another. The tree-spanning syntenic orthogroups are designated as phylogenetic anchors at this stage.

### V. Tandem Duplication Detection

To detect tandem duplicates, we implement another sliding window of size *D* genes. For each gene in a phylogenetic anchor group, we test whether the upstream or downstream *D* genes could be a tandem duplicate of the focal anchor gene (Fig. 5A and 5B). Let two organisms be denoted as species *A* and species *B*, their corresponding anchor genes be *a* and *b*, and a candidate duplicate as *a’*. To determine whether *a’* is a tandem duplicate of *a* without a direct comparison between the two, we examine the relationship between *a’* and *b*. We classify *a’* as a tandem duplicate if one of the three conditions are met. If *a’* has a BLAST hit to *b* and *b* is the top hit of *a’*, *a’* is flagged as a tandem duplicate (Fig. 5C). If this condition is not met, we compare the corrected bitscore between *a’* and *b*, labeled as *h’*, to the highest corrected bitscore found for *b* when comparing *A* and B, which we label as *h* (Fig. 5D). If *h’* is within *S* standard deviations (user defined) from *h* or if the two hits have the same e-value, *a’* is labeled as a tandem duplicate of *a*. We define this condition as the sequence similarity threshold condition. When neither of these conditions are satisfied, the neighboring *D* genes on both sides of *b* are scanned for genes that have their highest ranked BLAST hit directed towards *a’*, which we will denote as *b’*. If *b’* is already marked as a tandem duplicate to *b* or if *b’* has a BLAST hit to *a* and that corrected bitscore is within *S* standard deviations from the score between *b’* and *a’*, then *a’* is designated as a tandem duplicate to *a* (Fig. 5E). With the condition that a tandem duplicate is found, the search window is extended to the next *D* genes along the chromosome in the current window direction. A synopsis of the tandem duplication algorithm is illustrated in Figure 5.

**Figure 5:**
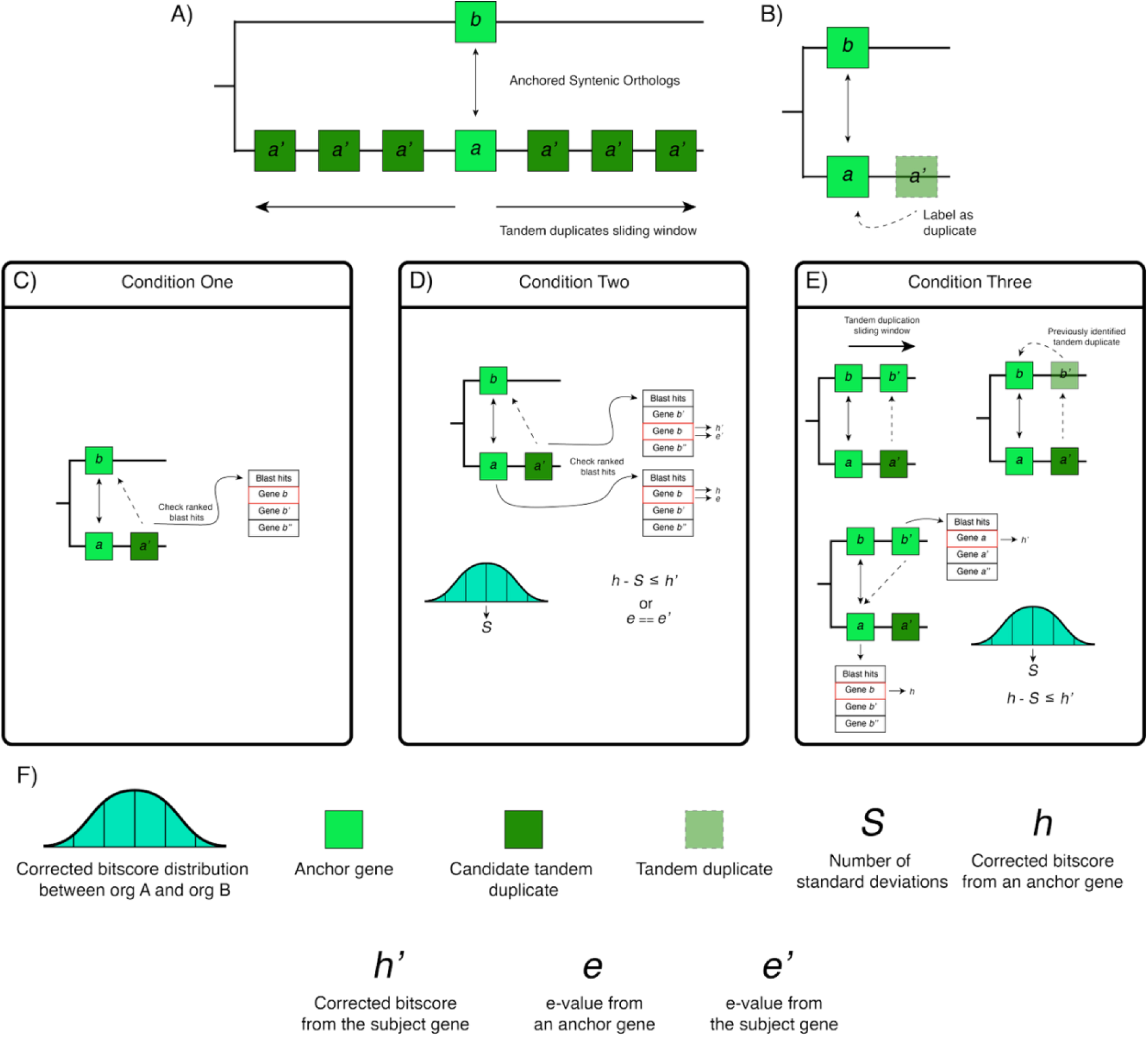
Summary of the tandem duplication algorithm. (A) Centering on an anchor gene, a sliding window is implemented in both the 3’ and 5’ directions to identify candidate tandem duplicates. (B) If a gene is marked as a tandem duplicate, it is merged to the anchor gene, effectively becoming invisible in the Synolog pipeline (C). A gene is marked as a tandem duplicate if it’s best BLAST hit is the interspecific anchor gene. (D) If the first condition is not met and if the candidate tandem duplicate has a BLAST hit to the interspecific anchor gene, that BLAST hit is compared to the focal anchor gene BLAST hit to determine if it is within the threshold. (E) In cases where the first two conditions are not met, if the candidate tandem duplicate has a match with a tandem duplicate of the interspecific anchor gene, and the previously labeled tandem duplicate’s best hit is the current anchor gene, the BLAST hits are again evaluated to determine tandem duplicate status. (F) Figures and symbols depicting the variables used in the tandem duplication algorithm are shown in below.

Under the scenario where no tandem duplicates were found for *A* when using *B* for the current phylogenetic anchor group, the group is queried for the next member down the guide tree, which we represent as *C*. The search for tandem duplicates is then repeated using ortholog *c* in place of *b*.

### VI. Updating Orthologs and Constructing Synteny Blocks

Before commencing with further ortholog detection using synteny, tandem duplicates that are not members of a phylogenetic anchor group are excluded from ortholog inference and synteny block construction. This tactic not only allows other genes to become orthologs, but it effectively makes these elements invisible to the sliding window applications. As such, a window size of *N* or *D* can cover more genomic space as the tandem duplicates would not contribute to these windows. Post filtering, the ortholog detection and synteny block construction follows that which is described in pipeline 1.

At the end of each iteration, new phylogenetic anchor groups are inferred without the requirement of enforcing the group to span the entire tree and for orthologs to contradict one another. For example, let’s consider a case with three organisms termed *A*, *B*, and *C* with homologs *a*, *b*, *b’*, and *c*. If *a* and *c* were considered orthologs, but were assigned to *b* and *b’* respectively, if *b* and *b’* were found to be co-localized within the same synteny blocks, the 5’ most duplicate would become the core phylogenetic anchor member while the other is flagged as a tandem duplicate. In the case where *a* is matched with *b*, *b* is matched with *c*, but no *C* ortholog was predicted for *a*, if there exist a bidirectional link between *a* and *c* from BLAST and these two genes are syntenic with *b*, then the phylogenetic anchor group can be assembled accordingly. Once no orthologs and tandem duplicates can be detected, the pairwise comparison ceases and the homolog relationships and synteny clusters are reported.

### VII. Orthogroup Characterization and Further Homolog Detection

After all the pairwise comparisons have been completed, phylogenetic anchor groups are merged if the constituting members were marked as a tandem duplicate but was assigned as a phylogenetic anchor beforehand. This merging process reconstructs a single phylogenetic anchor group with the core members consisting of the 5’ most gene. Each new phylogenetic anchor group is then scrutinized by determining the most basal organism in the group. Here, basal is defined as the organism closest to the root. This gene and the number of tandem duplicates is used to set the basal state for the orthogroup of interest. The remaining core members are then evaluated to assign the following states: if the gene count is greater than the basal number, a local expansion is assigned. When the gene count is lower than the basal number, a local contraction is granted to the homolog; otherwise, the gene retains a basal state.

Every orthogroup is examined to determine if every organism in the analysis is represented within the group. For every organism not in the orthogroup, each current member will be examined for a previously recorded ortholog from the missing organism established in the initial RBH approach. This backup ortholog is merged into the group if the ortholog is found to not be assigned to an orthogroup. In cases where multiple backup orthologs are found, the candidate orthologs are checked for shared synteny with the current core members and the first one meeting this criterion is selected. If no ortholog meets this condition, the first one reported is selected.

After updating the orthogroups, each core member in an orthogroup that lacks tandem duplicates is checked for additional tandem duplicates using the other core members as points of reference. This methodology serves to retrieve tandem duplicates for orthologs that were not anchor genes (i.e., not syntenic). If we call the current core member *a* from organism *A* and the current point of reference *b* from organism *B*, the search space is confined to all *A* genes matching *b* using the sequence similarity threshold conditional approach used in the standard tandem duplicate detection algorithm. *A* genes that are co-localized and are at most *D* genes away are merged as tandem duplicates. If a candidate duplicate is found to occupy a different chromosome and consist of a single exon, the candidate duplicate is merged into the orthogroup as a retrogene granted if *a* is composed of multiple exons.

### VIII. Segmental Duplication Detection

To detect segmental duplicates, we leverage the computed synteny blocks. The idea is to find sets of gene neighborhoods, currently without ortholog assignments, that resemble an established syntenic set of orthologs. If we label the two organisms constituting the synteny blocks *A* and *B*, we iterate over all of *A*’s genes in the synteny block and collect non-syntenic *A* genes that meet the sequence similarity threshold condition with the syntenic ortholog found in *B*. The collected genes are organized by chromosome and position, and a sliding window is used to find clusters of at least 3 genes that are no further apart than *S* genes from one another. Genes found within these clusters are tagged as segmentally duplicated genes. This procedure is then repeated for organism *B* with *A* being the point of reference. Once all the synteny blocks for *A* and *B* have been processed, the pairwise segmental duplications are reported.

Once all pairwise comparisons have reached completion, organism-specific segmental duplications are determined. For each organism, all tagged segmentally duplicated genes are re-organized by chromosome and position. A sliding window is then used to generate blocks of segmental duplications where the distance between any two genes in the segmental duplication block is no greater than the synteny sliding window size.

### IX. Tree Cluster Inference

Synolog is designed to identify synteny clusters that span across three or more organisms, which are termed tree clusters. The algorithm begins at the top (i.e., the tips) of the guide tree and initiates a syntenic ortholog search using synteny blocks between the current organism (*A*) and the next organism (*B*) down the guide tree. For each synteny block shared by *A* and *B*, each gene in *B* is queried for a syntenic ortholog found in *C*, where *C* is the next organism down the tree (i.e., the one sharing a most recent common ancestor) from *B* with a syntenic ortholog. If the tree clusters are to be constructed through a concurrent approach, the syntenic ortholog in *C* is required to be the syntenic ortholog with the current *A* ortholog. If the ortholog in *C* is validated, the *C* ortholog is added to the cluster and then used to get the next syntenic ortholog down the guide tree. This procedure is continued until there are less than three organisms that can be used to venture down the guide tree.

### X. Post Inference Toolkit

Synolog houses several utilities that allow the results of an analysis to be parsed or visualized. For instance, orthogroups containing a particular feature (e.g., orthogroups with local expansions for a specific species) can be readily extracted for evaluation using the `synolog_info.py` script. Visualizations can be generated from either a genome-wide, synteny block, regional, segmental duplication, or tree cluster perspective by invoking the `synolog_plot.py` subprogram. The `synolog_fasta.py` utility can be used to generate FASTA files containing gene sequences for orthogroups with the flexibility of querying only one-to-one orthogroups or requesting genes be represented by their longest transcript.

### Synteny-Based Scaffolding

The framework underpinning Synolog’s guided synteny scaffolding approach is designed to work in a pairwise fashion and is implemented in `synolog_collinearize.py`. The algorithm begins by a declaration of two genome assemblies: one to serve as a guide (*A*) and the other to be scaffolded (*B*). The conserved synteny blocks between two genome assemblies from running the Synolog pipeline in a prior analysis are then loaded (Fig. 6A). Contigs from *B* that contain regions of conserved synteny are bucketed, where each bucket represents the sequence in *A* that contained the most syntenic information for its corresponding assignees (Fig. 6B). Next, each synteny block is reduced to a pair of base pair positions that denote the midpoint positions of the synteny block in the respective sequences in *A* and *B*.

**Figure 6:**
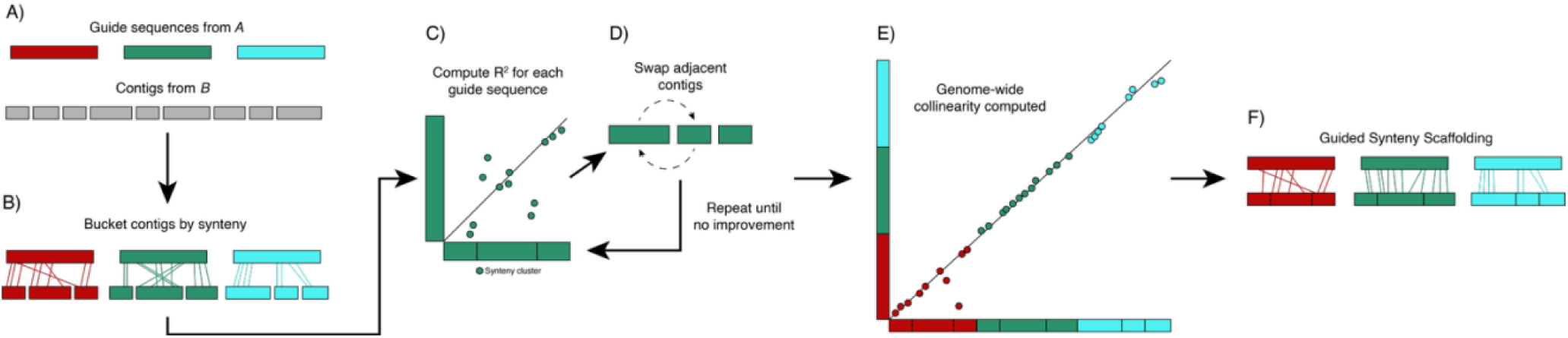
Outline of the synteny scaffolding algorithm. (A) The basis of synteny scaffolding algorithm begins with declaring a guide genome and a set of contigs to be scaffolded. (B) The contigs are bucketed to a guide sequence based on synteny content and then initially ordered based on the syntenic genes. (C) A hill-climbing algorithm is implemented where for each guide sequence, collinearity is measured using a linear regression with the contigs. (D) Contigs are swapped with adjacent neighbors after each iteration, with swaps that increase collinearity pursued. (E) After all guide sequences have a best fitted arrangement of contigs, a linear regression is performed using all guide sequences and the computed order of contigs. (F) This arrangement is then provided for users to use to scaffold the processed contigs.

To maximize the collinearity between the two genome assemblies, the sequences from *A* and *B* are projected onto two-dimensional space. Each sequence for *A* is assigned a fixed position while the contigs in *B* are repeatedly shuffled. To reduce the number of permutations, shuffling is performed within buckets and not between buckets. The initial permutation is constructed by ordering each sequence in *B* based on the average of adding each midpoint with respect to *A*. A linear regression is then computed between the chromosome in *A* and the contigs in *B* using the synteny midpoints, generating a R^2^ value (Fig. 6C). Next, adjacent sequences in *B* are swapped and a new R^2^ value is calculated and compared with the current score (Fig. 6D). The ordering with higher R^2^ score is retained as the current best and the analysis is repeated, with adjacent swaps performed and new orderings evaluated. If the score is worse than the current score, arrangements containing the offending swap are not pursued. The ordering with the highest R^2^ value is retained and designated as the best ordering of contigs for *B* for the given sequence in *A*.

After all sequences in *A* have been assigned a best ordering of sequences from *B*, a genome-wide linear regression is then performed between *A* and *B* (Fig. 6E). The set of instructions describing how the arrangement of contigs from *B* should be scaffolded is generated based on this genome-wide ordering and outputted by `synolog_collinearize.py`. Along with the targeted genome assembly and the respective annotations, these series of steps can be executed by `alter_genome_structure.py`, which will to perform the scaffolding and annotation lift over (Fig. 6F). An overview of the synteny scaffolding process can be viewed in Figure 6.

### Case Study 1: Testudine Evolution

To test the performance of Synolog in testudines, we obtained the reference genomes for the green sea turtle (Wang et al. 2013), the diamondback terrapin (Jiang et al. 2025), the red-eared slider (Simison et al. 2020), the Aldabra giant tortoise (Çilingir et al. 2022) and the Mexican gopher tortoise (http://www.vertebrategenomesproject.org/). Specific versions for each assembly are listed in Table S4. Non-coding genes were extracted from each assembly using the `synolog_ncgenes.py` utility and added to the species cache. All pairwise BLASTP and BLASTN analyses were executed employing the `synolog_blastctrl.py` tool for protein-coding and non-coding gene datasets, respectively. To arrange these species from an evolutionary perspective, we used the phylogeny reported by Thomson et al. (2021) to serve as the guide tree. The species were processed through the Synolog pipeline using the default values of a synteny sliding window of 100 genes, tandem duplication sliding window of 50 genes, and a standard deviation of 0.5 for the sequence similarity threshold condition.

To obtain gene symbols for gene ontology diagnosis, Synolog was re-executed including the Goode’s thornscrub tortoise (*Gopherus evgoodei*) (http://www.vertebrategenomesproject.org/) and Chicken (*Gallus gallus*) (https://useast.ensembl.org/Gallus_gallus/Info/Index?db=core) following pipeline 1 to obtain pairwise results. Genome versions for both assemblies can be found in Table S4. Each of the orthogroups of interest were assigned a single gene symbol from either Goode’s thornscrub tortoise or Chicken if the genes in the orthogroup did not have a single gene symbol. biomaRt v2.64.0 (Durinck et al. 2005) was used to query the gene symbols to the *G. evgoodei* gene ontologies, which were reduced using the R package rrvgo v1.20.0 (Sayols 2023) by applying the “Rel” method.

### Case Study 2: Metazoan Evolution

Echoing Simakov et al. (2022), the dataset consists of the genomes and annotations for Florida lancelet, the flame jellyfish, the Yesso scallop, the freshwater sponge *Ephydatia muelleri*, and the freshwater polyp. Except for the freshwater sponge and the Yesso scallop, the following accessions were obtained from NCBI. The genome for the freshwater sponge (Kenny et al. 2020) was downloaded from EphyBase (https://spaces.facsci.ualberta.ca/ephybase/) while the Yesso scallop dataset (Wang et al. 2017) was obtained from the MolluscDB (Liu et al. 2021, 2025). Specific genome accessions can be view in Table S5. For consistency, we followed the phylogeny presented in Simakov et al. (2022) to describe the evolutionary relationships between the five species.

The five species were processed through the Synolog pipeline similarly as with the first cast study with the exception of performing a more relaxed tandem duplication classification. Given the evolutionary distance amongst the samples, we ran Synolog and increased the standard deviation for the sequence similarity threshold condition from 0.5 to 2.0. Visualizations were generated using `synolog_ploy.py` for all syntenic genes and for tree-spanning genes, Lastly, to implement a similar method to that conducted by Simakov et al. (2022), we took the reported tree clusters produced by Synolog and searched for ALGs by merging tree clusters based on the encompassing chromosomes while ignoring genes on unplaced contigs or tree clusters lacking a member species. ALGs that only contained a single gene per species (i.e., did not contain any co-localized elements) were further excluded.

### Case Study 3: Synteny-Based Scaffolding

We obtained in-house contig-level reference genome for the mackerel icefish and the Antarctic bald notothen. The gene annotations for these assemblies were lifted over to the contig-based assembly using the respective AGP files and a custom python script. Following, we processed these contig-level assemblies through the Synolog pipeline using default parameters with the Patagonia blennie chromosome-level reference genome (Cheng et al. 2023). Evolutionary relationships were defined following the phylogeny predicted by Near et al. (2018).

The pairwise synteny information found with *E. maclovinus* was fed to `synolog_collinearize.py` for each of the two focal species while specifying *E. maclovinus* as the guide. The resulting organization, gene annotations, and contig-level reference genome was then processed by Synolog’s scaffolding submodule to generate a new annotated reference genome for both *C. gunnari* and *T. borchgrevinki*. The new assemblies were then added to the species cache and processed through the Synolog pipeline with the same *E. maclovinus* dataset. We then processed the two synteny-scaffolded genome assemblies though the `stats.sh` module from the BBTools suite (Bushnell 2014) to compare the assembly metrics to that presented in the publications.

### OrthoFinder2 Comparison

OrthoFinder2 is designed to work on protein sequences where a single isoform is used to represent each protein-coding gene. For case studies one and two, the longest transcript was selected for each gene using a custom python script. The filtered proteome was then processed through OrthoFinder2 v2.5.4 using default parameters. The resulting Phylogenetic Hierarchical Orthogroups (PHOs) were then compared to the orthogroups inferred by Synolog using a custom python script. Briefly, the orthogroups generated from Synolog are checked to determine if A) they are equivalent to any PHO, B) are a subset of a PHO, C) are a superset of a PHO, D) contain members in separate PHOs, or if they E) contain members not part of a PHO.

## Code Availability

Synolog (v1.0) is released under the free software, GPLv3 license. Source code can be downloaded from the Catchen lab website (http://catchenlab.life.illinois.edu/synolog/) and is made available in a public Git repository (https://bitbucket.org/jcatchen/synolog/src/master/). All datasets used in this study are publicly available in their respective studies.

## Acknowledgements

The authors would like to thank Malavika Venu, Angel G. Rivera-Colón for their helpful feedback when writing the manuscript, Katie Karl for her valuable advice regarding turtle biology, and Tatum Bernat and Andrew Whitehead for their input when testing the synteny-based scaffolding algorithm.

