## Supplemental Tables and Figures for "Synolog: A Scalable Synteny-Based Framework for Genome Architecture Characterization"

Table S1: Ortholog metrics from case study 1. The first column lists the species in this analysis, while the following two columns present the number of annotated protein-coding and non-coding genes. The number of protein-coding orthologs reported by both Synolog and OrthoFinder2 are then reported for each species. The number of genes in paralogous groups from OrthoFinder2 are reported for each species. The number of non-coding orthologs per species predicted from Synolog are shown.

| Species | Protein-Coding Genes | Non-Coding Genes | Synolog Protein-Coding Orthologs | OrthoFinder2 Orthologs | OrthoFinder2 Paralogs | Synolog Non-Coding Orthologs |
|---|---|---|---|---|---|---|
| *Chelonia mydas* | 19,937 | 8,554 | 19,174 | 19,007 | 452 | 2,353 |
| *Malaclemys terrapin centrata* | 21,126 | 5,109 | 20,009 | 20,125 | 607 | 2,357 |
| *Trachemys scripta elegans* | 18,785 | 3,671 | 18,223 | 18,243 | 239 | 1,973 |
| *Aldabrachelys gigantea* | 22,411 | 12,090 | 19,170 | 18,881 | 978 | 2,343 |
| *Gopherus flavomarginatus* | 21,033 | 7,988 | 19,370 | 19,279 | 1,262 | 2,699 |

Table S2: Ortholog metrics from case study 2. The species used in this study and number of annotated protein-coding genes are shown. The number of protein-coding orthologs reported by both Synolog and OrthoFinder2 are also presented. The last column shows the number of genes reported by OrthoFinder2 to be in lineage-specific orthogroups.

| Species | Protein-Coding Genes | Synolog Protein-Coding Orthologs | OrthoFinder2 Orthologs | OrthoFinder2 Paralogs |
|---|---|---|---|---|
| *Branchiostoma floridae* | 26,689 | 11,612 | 15,759 | 7,824 |
| *Rhopilema esculentum* | 14,865 | 10,325 | 11,053 | 2,267 |
| *Patinopecten yessoensis* | 24,733 | 11,341 | 11,540 | 4,367 |
| *Ephydatia muelleri* | 39,245 | 9,809 | 12,891 | 22,413 |
| *Hydra vulgaris* | 21,385 | 9,948 | 11,944 | 7,955 |

Table S3: Genome assembly metrics between synteny-based and Hi-C-based scaffolding assemblies from case study 3.

| | Synteny-Scaffolded L50 | Hi-C-Scaffolded L50 | Synteny-Scaffolded N50 | Hi-C-Scaffolded N50 |
|---|---|---|---|---|

| | | | | |
|---|---|---|---|---|
| *Champoscephalus gunnari* | 12 | 11 | 39.01 Mb | 44.08 Mb |
| *Trematomous borchgrevink* | 13 | 10 | 33.77 Mb | 42.66 Mb |

Table S4: Genome assemblies used in case study 1.

| Species | Common Name | Genome Version | Study |
|---|---|---|---|
| *Chelonia mydas* | Green Sea Turtle | GCF_015237465.2 | Wang et al. 2013 |
| *Malaclemys terrapin centrata* | Diamondback Terrapin | GCF_027887155.1 | Jiang et al. 2025 |
| *Trachemys scripta elegans* | Red-Eared Slider | GCF_013100865.1 | Simison et al. 2020 |
| *Aldabrachelys gigantea* | Aldabra Giant Tortoise | GCA_026122505.1 | Çilingir et al. 2022 |
| *Gopherus flavomarginatus* | Mexican Gopher Tortoise | GCF_025201925.1 | https://vertebrategenomesproject.org |
| *Gopherus evgoodei* | Goode's Thornscrub Tortoise | GCA_007399415.1 | http://www.vertebrategenomesproject.org/ |
| *Gallus gallus* | Chicken | GCA_016699485.1 | https://useast.ensembl.org/Gallus_gallus/Info/Index?db=core |

Table S5: Genome assemblies used in case study 2.

| Species | Common Name | Genome Version | Study |
|---|---|---|---|
| *Branchiostoma floridae* | Florida Lancelet | GCF_000003815.2 | Simakov et al. 2020 |
| *Rhopilema esculentum* | Flame Jellyfish | GCF_013076305.1 | Li et al. 2020 |
| *Patinopecten yessoensis* | Yesso Scallop | Patinopecten_yessoensis_hic_hwt_20211125 | Wang et al. 2017; Liu et al., 2021; 2025 |
| *Ephydatia muelleri* | Mueller's Freshwater Sponge | Emu_genome_v1 | Kenny et al. 2020 |
| *Hydra vulgaris* | Freshwater Polyp | GCF_022113875.1 | Simakov et al. 2022 |

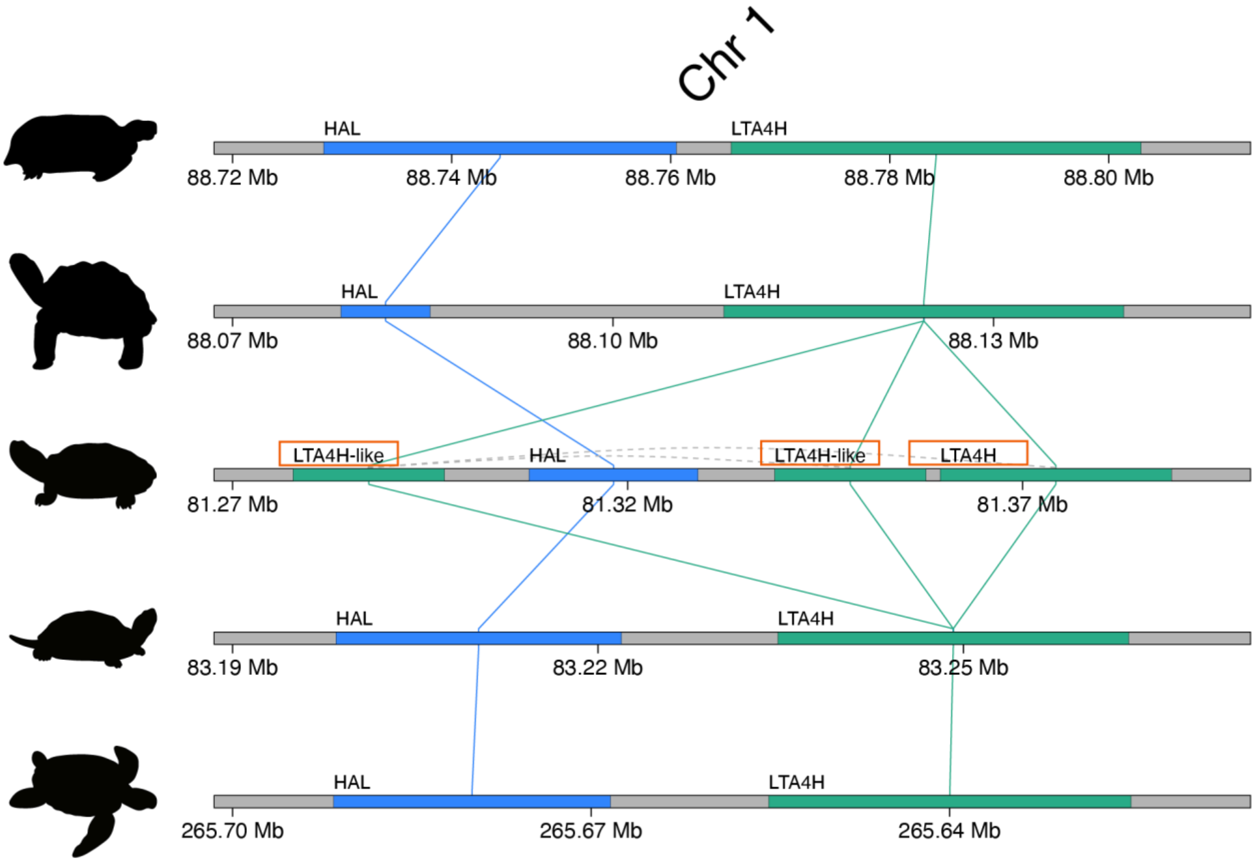


Figure S1: Tandem duplication of *LTA4H* in red-eared slider. Synolog reported a syntenic orthogroup of *LTA4H* where the red-eared slider was found to have three homologs. OrthoFinder2 (Emms and Kelly 2019) split this orthogroup into two groups, with the *LTA4H-like* genes encompassing a single paralogous group. Only syntenic genes were drawn.

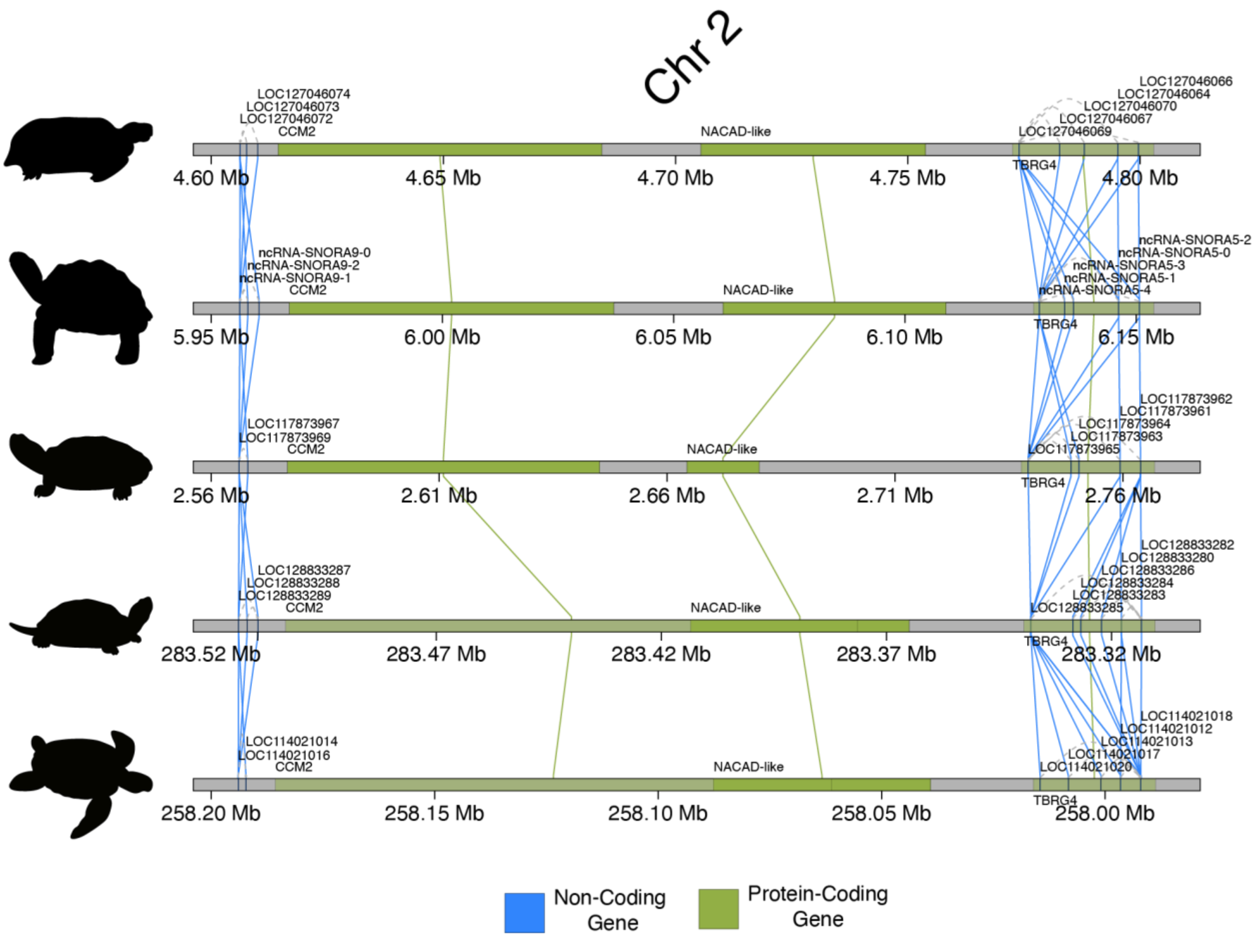


Figure S2: Example of syntenic non-coding orthologs in turtles. The syntenic region in chromosome 2 amongst turtles containing the *CMM2*, *NACAD-like*, and *TBRG4* genes habor putative syntenic non-coding genes that were automatically inferred by Synolog. Only syntenic genes are shown. Transparency for all overlapping genes was altered for clearer visualization.

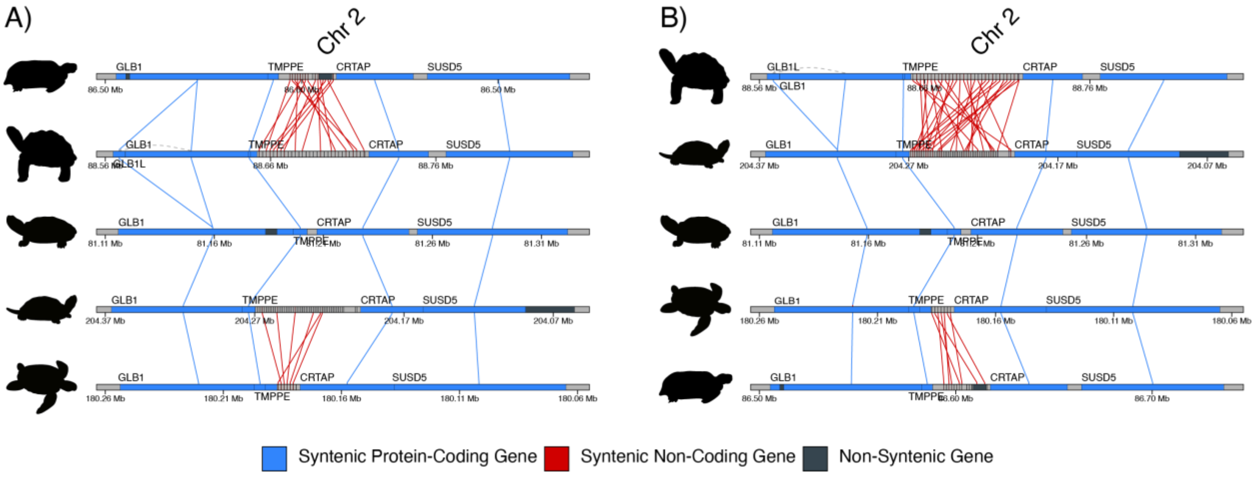


Figure S3: Illustration of a lineage-specific loss of non-coding genes in the red-eared slider. A) A locus containing tandem non-coding genes is found between the *TMPPE* and *CRTAP* genes for all species of turtles except for the red-eared slider. The ordering of the turtle species follows that used in the manuscript. B) Rearrangement of the species (from top to bottom, Aldabra giant tortoise, diamondback terrapin, red-eared slider, green sea turtle, Mexican gopher tortoise) highlights the varying syntenic content between the pairwise comparisons. Non-syntenic genes were all described as uncharacterized genes upon inspection.
